# Mitochondria-Associated Membranes (MAMs) are involved in Bax mitochondrial localization and cytochrome c release

**DOI:** 10.1101/443606

**Authors:** Alexandre Légiot, Claire Céré, Thibaud Dupoiron, Mohamed Kaabouni, Stéphen Manon

## Abstract

The distribution of the pro-apoptotic protein Bax in the outer mitochondrial membrane (OMM) is a central point of regulation of apoptosis. It is now widely recognized that parts of the endoplasmic reticulum (ER) are closely associated to the OMM, and are actively involved in different signalling processes. We adressed a possible role of these domains, called Mitochondria-Associated Membranes (MAMs) in Bax localization and fonction, by expressing the human protein in a yeast mutant deleted of MDM34, a ERMES component (ER-Mitochondria Encounter Structure). By affecting MAMs stability, the deletion of MDM34 altered Bax mitochondrial localization, and decreased its capacity to release cytochrome c. Furthermore, the deletion of MDM34 decreased the size of an uncompletely released, MAMs-associated pool of cytochrome c.

## Introduction

Apoptosis, the major programmed cell death pathway in animals, plays a central role during development, and along the whole life, by mediating the elimination of dispensable or potentially dangerous cells. Apoptosis is also involved in the response of cells to toxic molecules, such as anti-tumoral drugs. Apoptosis alterations are involved in developmental defects, tumor progression, and the resistance to anti-cancer treatments ([1,2] for reviews)

The permeabilization of the outer mitochondrial membrane (OMM) is a central event during the intrinsic pathway of apoptosis, that is normally activated by anti-tumoral treatments. It is controlled by proteins of the Bcl-2 family, among which the pro-apoptotic protein Bax is directly involved in the permeabilization process [3]. Following an apoptotic signal, Bax is redistributed from a mostly cytosolic to a mitochondrial localization that favors permeabilization. This redistribution is associated to the exposure of hydrophobic domains [4], the dimerization and the oligomerization of the protein [5], and changes in the interactions with different partners, including anti-apoptotic proteins of the Bcl-2 family, BH3-only proteins, and other regulators such as the mitochondrial receptor Tomm22 ([6-9] for reviews).

Mitochondria-Associated Membranes (MAMs) are a domain of the endoplasmic reticulum (ER) that is in close contact with the OMM [10,11]. They have been observed through microscopy and isolated as a membrane compartment containing elements from both ER and OMM. They are involved in biogenesis processes, like the biosynthesis and transfer of phospholipids from ER to mitochondria [12], in signaling processes, like ceramide synthesis [13] and Ca^2+^-movements between ER and mitochondria [14], or in degradation process, like mitophagy [15]. MAMs are dynamic structures that are done and undone, depending on the physiological status of the cell ([16-18] for reviews).

We addressed a possible role of MAMs in Bax localization and redistribution in healthy cells and during apoptosis. Under certain conditions, a small proportion of Bax was found in ER [19,20]. The hypothesis that MAMs stability could contribute to the regulation of Bax movements between different compartments thus deserves investigations. As underlined above, a difficulty comes from the fact that MAMs are not a stable well-defined compartment but are dependent on other cellular processes such as mitochondrial dynamics, Ca^2+^-signaling, or mitophagy. In mammalian cells, it is then difficult to evaluate the contribution of MAMs in Bax localization independently from these cellular processes, that also modulate apoptosis.

The heterologous expression of Bax, and other Bcl-2 family members, in yeast is a simplified cellular model tp investigate molecular mechanisms underlying the interaction of Bax with mitochondria [21]. In yeast, MAMs stability depends on the ERMES complex (ER-Mitochondria Encounter Structure). ERMES is formed of 4 proteins, Mmm1p, Mdm10p, Mdm12p and Mdm34p ([22],[17] for review). A fifth protein, Gem1p, belonging to the Miro family (Mitochondria Rho GTPase) contributes to the regulation of the complex [23]. The role of ERMES in lipid transfer between ER and OMM has been largely documented ([16] for review), but remains discussed [24]. Also, the role of individual proteins remains unclear. Furthermore, the deletion of MMM1, MDM10 or MDM12 genes leads to a dramatically altered phenotype, including a rapid loss of mitochondrial DNA, rendering functionnal studies difficult [25,26]. However, the deletion of MDM34 leads to a marginally altered phenotype, with functional mitochondria (this study). We thus expressed Bax in a MDM34-null mutant, to evaluate the dependence of Bax localization and function on MAMs stability. Our results show that Bax mitochondrial localization is decreased when MAMs were destabilized through the deletion of MDM34. Furthermore, we observed a pool of incompletely released, MAMs-associated cytochrome c, the size of which was decreased in the *Δmdm34* strain.

## Experimental procedures

The deletion cassettes of ERMES genes from the Euroscarf collection were amplified by PCR, transferred into the W303-1A strain by homologous recombination and selection for G418 resistance, and verified by PCR with primers located within the KanMX4 gene and in the 5’ and 3’ sequences of ERMES gene. Like reported by others [26], the deletion of MMM1, MDM10 or MDM12 led to a rapid loss of mitochondrial DNA (*rho-/0* genotype), making these strains useless for our studies. However, we could generate stable *rho+* strains deleted for MDM34.

Wild type W303-1A (*mat*a, *ade2*, *his3*, *leu2*, *trp1*, *ura3*) and the mutant *Δmdm34*::kanMX4 were transformed with plasmids pYES3-BaxWT, pYES3-BaxP168A, or pYES3-BaxS184V [27,28]. Co-transformations with the plasmid pYES2-Bcl-xL were also done.

Cells were pre-grown in YNB medium (Yeast Nitrogen Base 0.17%, ammonium sulfate 0.5%, potassium dihydrogenphosphate 0.1%, Drop-Mix 0.2%, auxotrophic requirements 0.01%, pH 5.5, supplemented with 2% glucose, and then transferred in the same YNB medium supplemented with 2% lactate to obtain an optimal differentiation of mitochondria. When the cultures reached the mid-exponential growth phase (O.D. at 550nm between 4 and 6), they were diluted in the same medium down to 0.7 - 0.8 O.D. and added with 0.8% galactose, to induce Bax expression.

Whole lysates were prepared from 10-20 mL cultures. Bax expression was induced for 5 hours. Cells were harvested and washed in the RB buffer (0.6M mannitol, 2mM EGTA, 10mM tris-maleate 10mM, pH 6.7, anti-proteases cocktail (Complete-Mini, Roche)), resuspended in 0.5mL of ice-cold RB buffer and added with 0.40mm-mesh HCl-washed glass beads (2/3 of the total volume). Cells were broken in a Tissue Lyser (30Hz, 3 minutes). The homogenate was centrifuged at 500 x g (10 minutes) to eliminate residual beads, unbroken cells and nuclei. Protein concentration was measured by the Lowry method.

Crude mitochondria fractions were isolated from 2L-cultures. Bax was expressed for 14 hours (see [28] for a detailed protocol). Briefly, cells were converted to spheroplasts and homogeneized. Two series of low-speed/high-speed centrifugations allowed to obtain a crude mitochondria pellet.

1-3 mg of proteins from cell lysate or isolated mitochondria (10mg/mL) were layered on the top of a 10 mL-density gradient in Beckmann SW41 tubes. Both sucrose and Optiprep (Stemcell Technologies) gradients were done (see results). Gradients were centrifuged overnight at 135,000 x g (28,000 rpm). Fractions were recovered with an Auto-densiflow II collector (Buchler), precipitated with 0.3M TCA, washed with 100μL cold acetone and solubilized in Laemmli buffer.

Cytochromes contents of isolated mitochondria were measured by differential redox spectrophotometry in a double-beam spectrophotometer Varian Cary 4000, as described previously [29].

For western-blots, proteins were separated on 12.5% SDS-PAGE, transferred onto nitrocellulose, saturated in PBST/milk or TBST/BSA (depending on antibodies) and blotted with the following antibodies: anti-human Bax 2D2 (Santa-Cruz, 1/5,000 dilution), anti-yeast porin (Por1) (Novex, 1/50,000 dilution), anti-yeast Phosphoglycerate Kinase (Pgk1) (Novex, 1/10,000 dilution), anti-yeast Dolichol Phosphate Mannose Synthase (Dpm1) (Novex, 1/5,000 dilution), anti-yeast Cytochrome c (Cyc1) (Custom made, Millegen, 1/5,000 dilution), anti-yeast Cytochrome c Oxidase subunit II (Cox2) (Invitrogen, 1/5,000 dilution, anti-yeast carboxypeptidase Y (Cpy1) (Invitrogen, 1/10,000 dilution). Horse radish peroxidase-coupled secondary antibodies (Jackson Laboratories, 1/10,000 dilution) were revealed by ECL (Luminata Forte, Millipore), vizualized with a digital camera (G-Box, Syngene), and quantified using Image J software. Statistical analyzes and figures were done with GraphPad Prism 6 software.

## Results and discussion

A substitution P168A stimulates Bax mitochondrial localization and ability to permeabilize the OMM to cytochrome c when expressed in yeast [27,28] and in glioblastoma cells [30]. Also, purified recombinant Bax-P168A was more efficient than the wild-type protein to permeabilize isolated yeast or human mitochondria [31]. The difference between the two proteins was largely attenuated when assayed on liposomes [31] suggesting that additional components were needed to emphasize the distinct activities of the two proteins. Interestingly, a mutant P168G forms stable dimers not inserted into the mitochondrial membrane [32]. This substitution does not simply change the position of the hydrophobic helix α9, but has more general effects on the whole conformation of Bax [33]. Considering these data, we studied whether other factors that have not been considered until now could be involved in the ability of Bax-P168A to interact with mitochondria.

We built a W303-1B strain carrying a *Δmdm34::kanMX4* deletion (that will be named *Δmdm34* below). This strain did not exhibit major alterations of cell growth and viability, opposite to mutants carrying deletions of MMM1, MDM10 or MDM12 [25,26;34-37]. Indeed, *Δmdm34* grew in YNB supplemented with the non-fermentable carbon source lactate, showing that mitochondria were functionnal. Compared to wild-type, the doubling time was marginally increased (4h15 *v/s* 4h) and the cell density at the stationnary phase was slightly decreased (2×10^8^ *v/s* 2.4×10^8^ cells/mL). This suggested that the deletion of MDM34 did not impair the replication/transmission of mitochondrial DNA.

Cellular extracts of the W303-1B and *Δmdm34* strains were analyzed on Optiprep density gradients (Fig.1). Differences between the two strains were in line with the expected phenotype of *Δmdm34*. In wild-type extracts, a minor population of ER (vizualized by the presence of Dpm1) was found at a higher density (around fraction 12) than the main ER population (fractions 3-8). This fraction was absent from *Δmdm34* extracts. However, fractions containing mitochondrial proteins (Por1 and Cox2) were found at lower densities in *Δmdm34* (fractions 10-17) than in wild-type (12-17). Consequently, the *Δmdm34* strain still contained fractions where ER and mitochondrial proteins were both present (namely fraction 10), albeit at a lower density than in wild-type (fractions 12-13).

**Figure 1:**
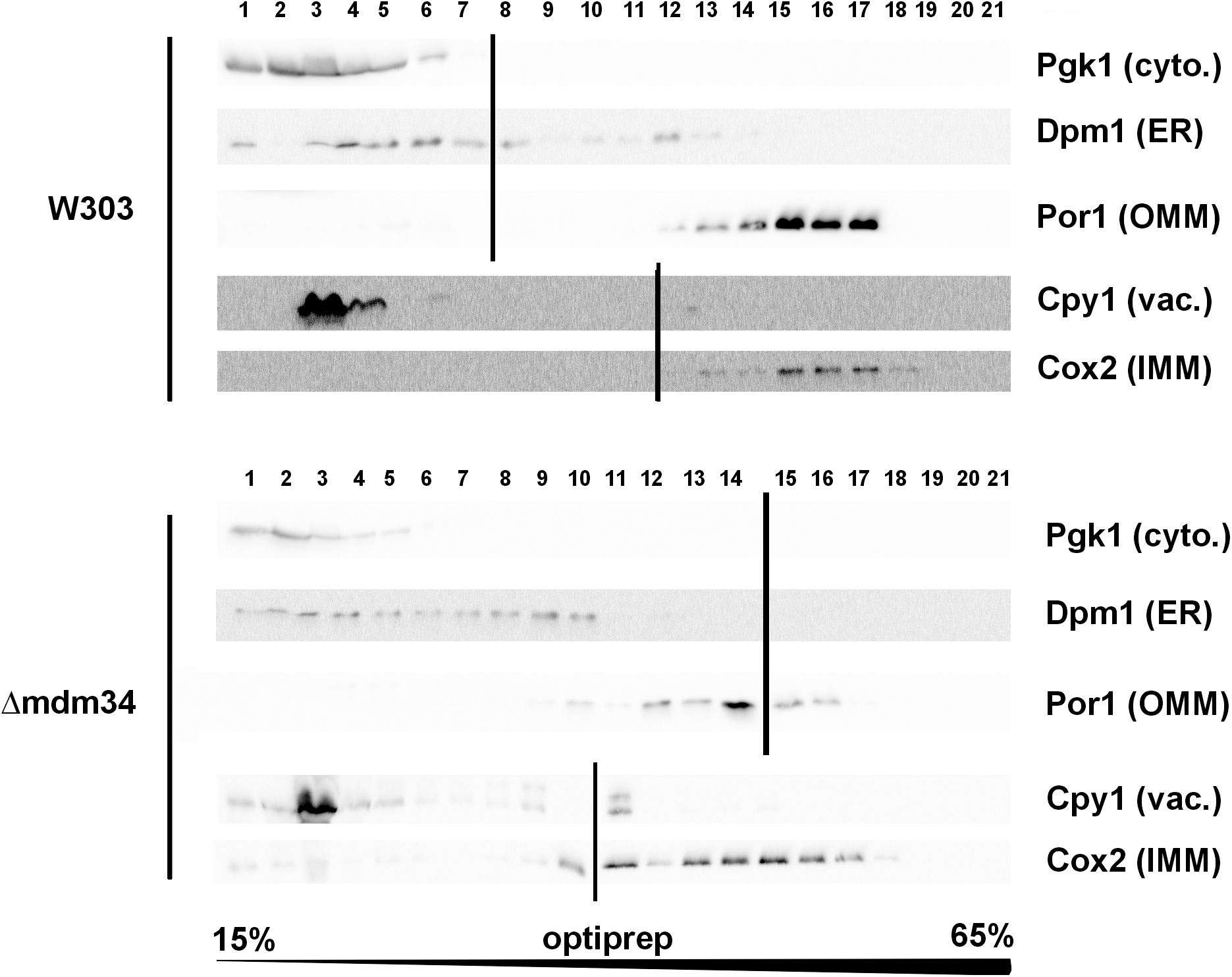
The deletion of MDM34 destabilizes MAMs. Cell extracts were separated on a 15-65% gradient of Optiprep. The gradient was resolved into 21 fractions and analyzed by western-blot for the presence of indicated markers. Vertical lines separate different gels. Data shown are from extracts obtained at the same time for each strain, and are representative of four independent experiments.

To test the involvement of these modifications on Bax relocation and activity, the mutant Bax-P168A was comparatively expressed in wild-type and *Δmdm34* yeast strains. In yeast, the activity of Bax is quantitatively measured through the amount of residual mitochondrial cytochrome c by redox spectrophotometry: less mitochondrial cytochrome c indicates a higher Bax activity. The activity of Bax-P168A was slightly but significantly decreased when expressed in *Δmdm34* strain (Fig.2A). The coexpression of the anti-apoptotic protein Bcl-xL partially inhibited Bax-P168A-induced cytochrome c release (Fig.2B), and abolished the difference between both strains, showing that the initial difference actually reflected an alteration of Bax-P168A activity and not a general property of *Δmdm34* mitochondria. Cell extracts from both strains expressing Bax-P168A were analyzed on sucrose density gradients. Although sucrose-based gradients showed more cross-contamination than Optiprep-based gradients, they provided a better discrimination between ER-containing fractions in wild-type and *Δmdm34* strains (Fig.2C). Under these conditions, a population of Bax-P168A was associated to ER in *Δmdm34* strain (fraction 4) but not in wild-type (Fig.2C). This suggested that some Bax-P168A remained associated to the ER compartment, resulting in the decreased ability to release cytochrome c.

**Figure 2:**
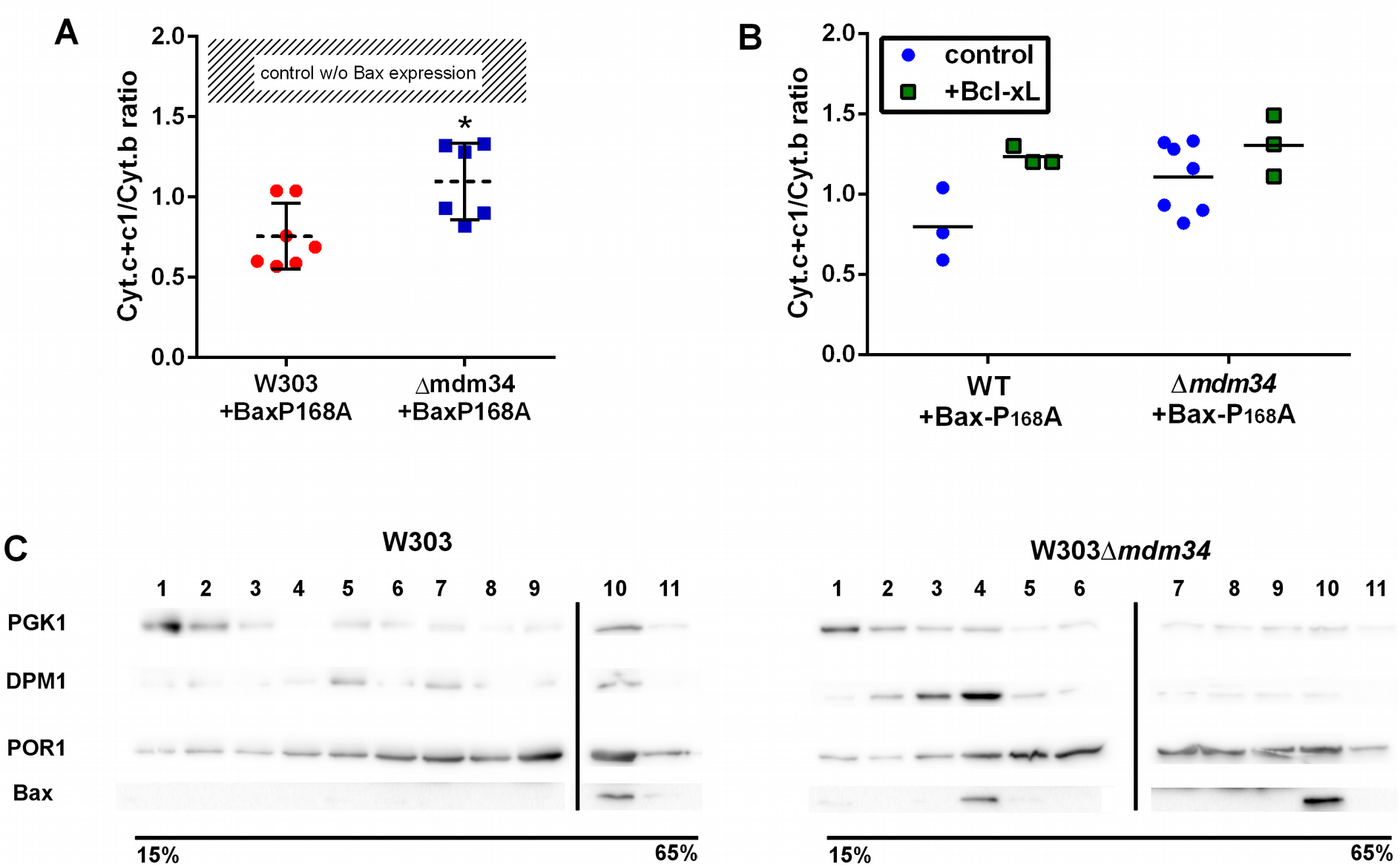
Bax-P168A is partly retained in ER, and has a decreased ability to release cytochrome c in *Δmdm34* cells. (A) Cytochrome c+c1/Cytochrome b ratios measured on mitochondria isolated from wild-type or *Δmdm34* cells expressing Bax-P168A. Each point represents a single mitochondria preparation. The hatched zone corresponds to the typical values found on mitochondria preparations from cells that do not express Bax [27-39]. *: p<0.05 (unpaired Student t-test). (B) Cytochrome c+c1/Cytochrome b ratios measured on mitochondria preparations isolated from strains co-expressing Bax-P168A and Bcl-xL. (C) Separation of whole extracts from cells expressing Bax-P168A on a 15-65% sucrose density gradient. Vertical lines mark the separation between different gels. Data are representative of four independent experiments.

Ser184 is located in the helix α9 of Bax, and is the target of several kinases, including AKT [38,39]. The phosphorylation of Bax by AKT impairs Bax action on mitochondria, but the underlying mechanisms are not completely clear, as they depend on the presence of anti-apoptotic proteins [40,41]. However, there is a consensus about the consequences of substitutions of this residue on Bax localization: Bax-S184D remains mostly cytosolic (and is rapidly degraded when expressed in yeast), while Bax-S184V (or Bax-S184A) is markedly associated to mitochondria (including in yeast).

Although being largely mitochondrial, Bax-S184V was less efficient than Bax-P168A to promote the release of cytochrome c, like previously reported [40]. Furthermore, it was less efficient when expressed in *Δmdm34* strain than in the wild-type strain, suggesting that the *Δmdm34* deletion had a stronger effect on Bax-S184V than on Bax-P168A (Fig.3A). As negative controls, *Δmdm34* strain that did not express Bax, or that expressed BaxWT (which, like in non-apoptotic mammalian cells, is poorly mitochondrial), did not show any difference in residual mitochondrial cytochrome c, in comparison to wild-type.

**Figure 3:**
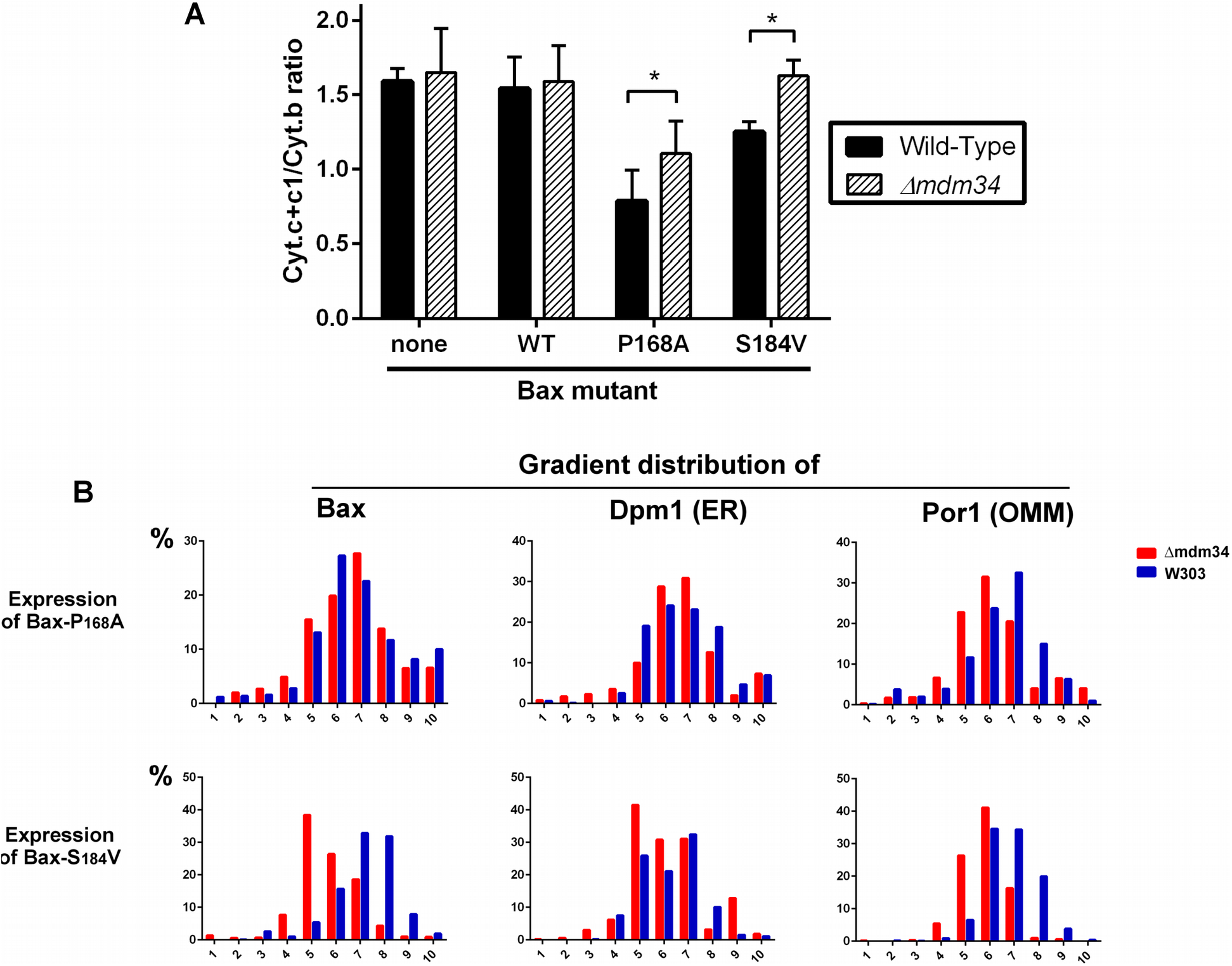
Bax-S184V looses the ability to release cytochrome c in *Δmdm34* cells. (A) Cytochrome c+c1/Cytochrome b ratios measured on mitochondria from different strains. Values are averages (± s.d.) of 4 to 6 independent experiments. (B) Distribution of Dpm1 (ER), Por1 (OMM) and Bax on a crude mitochondrial fraction separated on sucrose density gradients. Results are representative of 3 independent experiments.

We next analyzed the distribution of crude mitochondria preparations of strains expressing Bax-P168A and Bax-S184V on density gradients (Fig.3B). Like for gradients of whole cells extracts, we observed that mitochondria-rich fractions (containing Por1), were less dense in *Δmdm34* strain (around fraction 6) than in the wild-type strain (around fraction 7).

The distribution of Bax-P168A was not dramatically altered by the *Δmdm34* mutation, with a marginal inversion between the two major Bax-containing fractions (6 and 7). The *Δmdm34* mutation slightly retained Bax-P168A in the ER (Fig.2C and 3B) but enough protein remained able to reach the mitochondria to promote a significant release of cytochrome c, although slightly lower than in the wild-type strain (Fig.3A).

The change in the distribution of Bax-S184V was more dramatic, with a marked shift towards lower densities fractions (5 and 6). It is noteworthy that fraction 5, that contained 40% of total Bax-S184V present in the crude fraction, contained 40% of total Dpm1, but less than 30% of Por1, suggesting that a significant proportion of Bax-S184V remained localized in ER membranes associated to the crude mitochondrial fraction: this correlates with the total loss of capacity of Bax-S184V to release cytochrome c in *Δmdm34* strain (Fig.3A).

Experiments reported above showed that MAMs stability affect, albeit moderatly, the mitochondrial localization of Bax and its capacity to release cytochrome c. We next adressed whether cytochrome c release was, by itself, affected by MAMs stability. Our hypothesis was that, following Bax expression, the mitochondrial fraction might contain two pools of cytochrome c: unreleased cytochrome c genuinely localized in the intermembrane space or ‘uncompletely released’ cytochrome c that was not anymore in the intermembrane space but not yet in the cytosol, and that was trapped in ER membranes associated to mitochondria. To discriminate between these pools, we compared the level of reduction by dithionite, a chemical reducer, with the level of reduction by NADH, a substrate of the yeast respiratory chain, that is oxidized by two intermembrane space-facing NADH dehydrogenases, thus able to reduce only cytochrome c localized within the intermembrane space [42].

On wild-type mitochondria, as expected, 100% of cytochrome c reduced by dithionite was also reduced by NADH (Fig.4A). However, on mitochondria isolated from the strain expressing Bax-P168A, a significant proportion of cytochrome c reduced by dithionite was not reduced by NADH (Fig.4A). This showed the existence of a pool of cytochrome c, still present in the mitochondrial fraction but not in the intermembrane space, thus not able to receive electrons transferred from Complex III. We next tested whether there was a difference between wild-type and *Δmdm34* strains. To attenuate the effect of variations in purity between mitochondrial preparations, we corrected the reduction of cytochrome c by the reduction of cytochrome b, an unreleased integral membrane protein (Fig.4B). There was no difference between wild-type and *Δmdm34* control strains, showing that, in the absence of Bax, all cytochrome c remained within the intermembrane space. There was also no difference between wild-type and *Δmdm34* strains expressing Bax-P168A. As reported above (Fig.4A), a fraction of cytochrome c was not reducible by NADH: Fig.4B showed that the size of this fraction was the same in wild-type and *Δmdm34.* We found a different behaviour in strains expressing Bax-S184V: the size of the NADH-reducible pool was decreased in the wild-type, and significantly re-increased in *Δmdm34* (Fig.4B). This showed that a pool of non NADH-reducible cytochrome c remained in the mitochondrial fraction following the expression of Bax-S184V, and disappeared in the *Δmdm34* strain: this uncompletely released pool of cytochrome c was thus dependent on MAMs stability. Supporting this hypothesis, in mitochondria preparations from wild-type cells expressing Bax-S184V, but not in *Δmdm34* expressing Bax-S184V, some cytochrome c was found in lower density fractions, that may correspond to the partly or incompletely released cytochrome c identified by spectrometry (Fig.4C).

**Figure 4.**
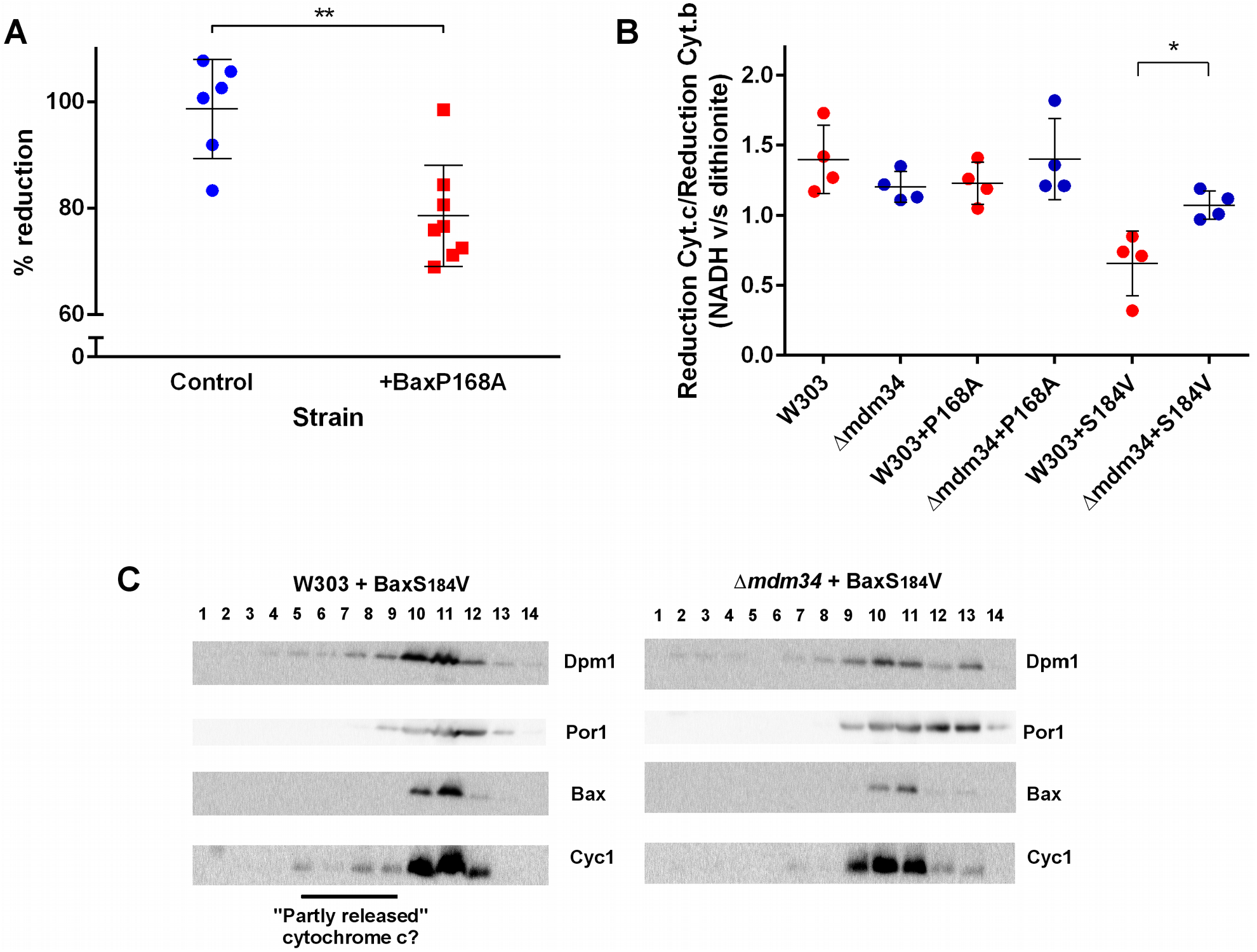
An incompletely released, MAMs-associated, pool of cytochrome c is decreased in *Δmdm34* cells. (A) The fraction of NADH-reducible cytochrome c was measured by measuring reduction after adding 1mM NADH, followed by measuring the reduction after adding dithionite on the same sample. Each point is a different mitochondria preparation (**: significantly different p<0.01). (B) The ratio reduction by NADH/reduction by dithionite was measured for cytochrome c and for cytochrome b. The graph shows the ratio (% reduction cytochrome c) / (% reduction of cytochrome b). Each point is a different mitochondria preparation. (*: significantly different p<0.05). (C) Mitochondria were fractionated on a sucrose density gradient. A small pool of cytochrome c was found at a lower density than the main fraction, that may correspond to partly released cytochrome c, not reducible by NADH in experiments (A) and (B). The figure is representative of 3 independent experiments.

Data reported above show that, in a model cellular system, the interaction of Bax with mitochondria is modulated by the stability of MAMs. The effects of MDM34 deletion on MAMs stability only marginally affected cell growth and viability, contrary to the deletion of the three other ERMES components. This is in line with the proposed model for ERMES organization, where the absence of MDM34 would have less dramatic effects on the complex organization than the absence of any of the three other ones [17]. In spite of this moderate effect on MAMs stability, MDM34 deletion was sufficient to alter the behavior of both Bax-P168A and Bax-S184V mutants.

Both mutants display a strong mitochondrial localization, but not the same level of activity. Bax-P168A is very active, as evidenced by a high ratio of cytochrome c release [28,30,31]. On the opposite, the activity of Bax-S184V is modest [28,40], particularly in regard to its high mitochondrial content, suggesting that its intrinsic activity is low [29,40]. The consequences of MDM34 deletion were thus more visible on Bax-S184V than on Bax-P168A, both in terms of Bax localization and of capacity to release cytochrome c. This revealed an unexpected aspect of Bax function: a fraction of cytochrome c that was still present in the crude mitochondrial fraction, could not be reduced by the respiratory substrate NADH. This pool of cytochrome c was detected by spectrophotometry, showing its full assembly with the haem moiety, a process taking place within mitochondria ([43] for review), and eliminating the hypothesis that it was a pool of cytochrome c not reaching mitochondria. It was therefore a pool of released cytochrome c, that remained yet associated to mitochondria. This fraction of cytochrome c likely corresponds to that was detected by western-blots in lower density fractions. Interestingly, both NADH reduction and western-blots showed that the size of this fraction was markedly decreased in *Δmdm34*, evidencing that this fraction of incompletely released cytochrome c was actually located in MAMs.

These data show that, in a model cellular system, both Bax relocation to mitochondria and cytochrome c release to the cytosol depend on the stability of MAMs, suggesting that this compartment may play a role in the regulation of Bax activity during apoptosis. These observations should now be extended to mammalian cells where it is, however, more difficult to modulate MAMs stability without affecting other processes relevant to apoptosis.

## Aknowledgements

This work were supported by the Centre National de la Recherche Scientifique and the Université de Bordeaux.

